# Normative brain-state trajectories reveal deviation from healthy aging in AD

**DOI:** 10.64898/2026.06.19.733190

**Authors:** Monireh Taimouri, Vikram Ravindra

## Abstract

**INTRODUCTION:** Distinguishing healthy brain aging from early neurodegenerative disruption remains a major challenge in Alzheimer’s disease.

**METHODS:** Using resting-state fMRI from the Alzheimer’s Disease Neuroimaging Initiative (357 participants; cognitively normal, mild cognitive impairment, Alzheimer’s disease), we trained a Hidden Markov Model exclusively on cognitively normal adults to define latent connectivity states. A generalized additive model estimated an age-adjusted reference trajectory of transition entropy, and subject-specific deviations from this trajectory were quantified.

**RESULTS:** Mild cognitive impairment and Alzheimer’s disease showed progressively greater deviation from the normative reference, with the strongest disruption in Alzheimer’s disease. A single absolute-deviation score retained much of the predictive information contained in higher-dimensional dynamic features.

**DISCUSSION:** These findings suggest Alzheimer’s disease is associated with measurable departure from healthy dynamic brain aging, providing an interpretable framework for future longitudinal and clinically validated studies.

## 1 Introduction

Aging is accompanied by substantial heterogeneity in brain structure, function, and cognitive resilience. A central challenge in aging research is therefore to distinguish variation that reflects normative aging from deviations that signal emerging age-related disease. This distinction is especially important for Alzheimer’s disease (AD), where pathological changes unfold against a background of broad inter-individual variability in healthy older adults [1–4]. As a result, many neuroimaging biomarkers struggle to determine whether an observed pattern reflects disease, normal aging, or simply one person’s position within the broad range of healthy variation. This problem is especially acute in the prodromal stages of dementia, where subtle abnormalities may already be present but remain difficult to separate from normative aging effects [5–7].

Most existing functional MRI biomarkers rely on static summaries of connectivity, implicitly treating brain networks as temporally stable entities. However, brain function is inherently dynamic: even during rest, large-scale networks recurrently reconfigure over time, alternating between transient but structured patterns of interaction [8–10]. These fluctuations are increasingly understood as a fundamental property of healthy brain organization, linked to cognition, flexibility, and large-scale coordination [11, 12]. This raises a compelling possibility for neurodegeneration: the most informative signal of pathology may lie not only in *which* networks are connected, but in *how* the brain moves between connectivity states over time [6, 13, 14].

State-based modeling approaches provide a natural framework for capturing this temporal organization. In particular, Hidden Markov Models (HMMs) offer a probabilistic description of brain dynamics as transitions between latent connectivity states, each associated with a characteristic pattern of network interactions. Applied to resting-state fMRI, HMMs yield interpretable dynamic biomarkers, including how frequently particular states are occupied, how persistently they are maintained, and how predictably transitions occur between them. Prior work has demonstrated that HMMs can reveal reproducible and hierarchically organized large-scale brain states, establishing them as a principled representation of the dynamic architecture of brain function [12, 15, 16].

Yet most studies of dynamic functional connectivity in aging and AD remain limited in two important respects. First, they typically rely on group-level comparisons, which reveal average disease effects but do not indicate whether a given individual’s dynamics are abnormal *for their age*. Second, even when dynamic features are extracted, they are usually treated as fixed endpoints rather than quantities that can be embedded within a normative aging trajectory. Consequently, current approaches provide limited insight into whether altered brain dynamics in MCI or AD represent a continuation of healthy aging or a genuine deviation from it [6, 17–19].

Normative modeling offers a way forward. By learning age-dependent trajectories from healthy individuals and quantifying how far new observations deviate from those trajectories, normative models move beyond case-control averages toward individualized estimates of abnormality. This strategy has proven useful across neuroimaging settings, but has rarely been extended to dynamic functional biomarkers and has not been systematically integrated with HMM-based descriptions of brain-state dynamics in the context of aging and Alzheimer’s disease [18–21].

Here, we propose a normative dynamic-state framework for characterizing large-scale brain dynamics across the continuum from healthy aging to mild cognitive impairment (MCI) and AD. Using resting-state fMRI from the Alzheimer’s Disease Neuroimaging Initiative (ADNI), we first estimate dynamic functional connectivity and represent it in a low-dimensional latent state space using a Hidden Markov Model trained on cognitively normal individuals only. We then model age-adjusted normative trajectories of transition entropy using generalized additive models, and quantify subject-specific deviation from this healthy reference. This framework allows us to ask a more biologically precise question than standard classification: not simply whether clinical groups differ, but whether brain-state dynamics show progressively greater disruption relative to an age-adjusted healthy baseline.

We show that dynamic biomarkers derived from this framework capture an ordered shift from cognitively normal aging to MCI and then to AD. In particular, transition entropy and deviation from the normative trajectory identify abnormal disruption of latent brain-state organization, with the clearest effects observed in AD and intermediate alterations in MCI. Together, these results suggest that deviation from normative brain dynamics can provide a compact and interpretable candidate biomarker of disease-related disruption in older adults.

## 2 Results

We first defined an age-adjusted reference model of brain-state dynamics in cognitively normal older adults, and then used it to quantify disease-related deviations in MCI and AD.

### 2.1 An age-adjusted reference model captures normative brain-state dynamics

To characterize normative age-related changes in large-scale brain dynamics, we first modeled the relationship between age and metastability using cognitively normal (CN) subjects only. Metastability was quantified as the transition entropy of subject-specific HMM state sequences, capturing the degree of randomness in transitions between latent functional connectivity states.

Within each cross-validation fold, an age-adjusted generalized additive model (GAM) with a smooth term for age (up to 10 splines) was fitted using training CN subjects only. The fitted model was then applied to held-out CN, MCI, and AD subjects to estimate the expected entropy under normative aging.

Figure 1 compares a descriptive GAM fitted to all CN scans with a GAM fitted to held-out CN scans only. The two curves were broadly similar across the observed age range, with a mean absolute difference of 0.019 and a maximum absolute difference of 0.074, indicating that the normative CN trajectory was reasonably stable under held-out evaluation. Transition entropy in held-out CN subjects remained low overall (mean = 0.113, SD = 0.139), consistent with only modest age-related variation. In a descriptive GAM fitted to all CN scans, the smooth age term was statistically significant (*p* = 2.70 × 10^−3^), but the explained deviance was small (pseudo-*R*^2^ = 0.023), indicating that age accounted for only a limited fraction of the variability in transition entropy.

**Fig. 1.**
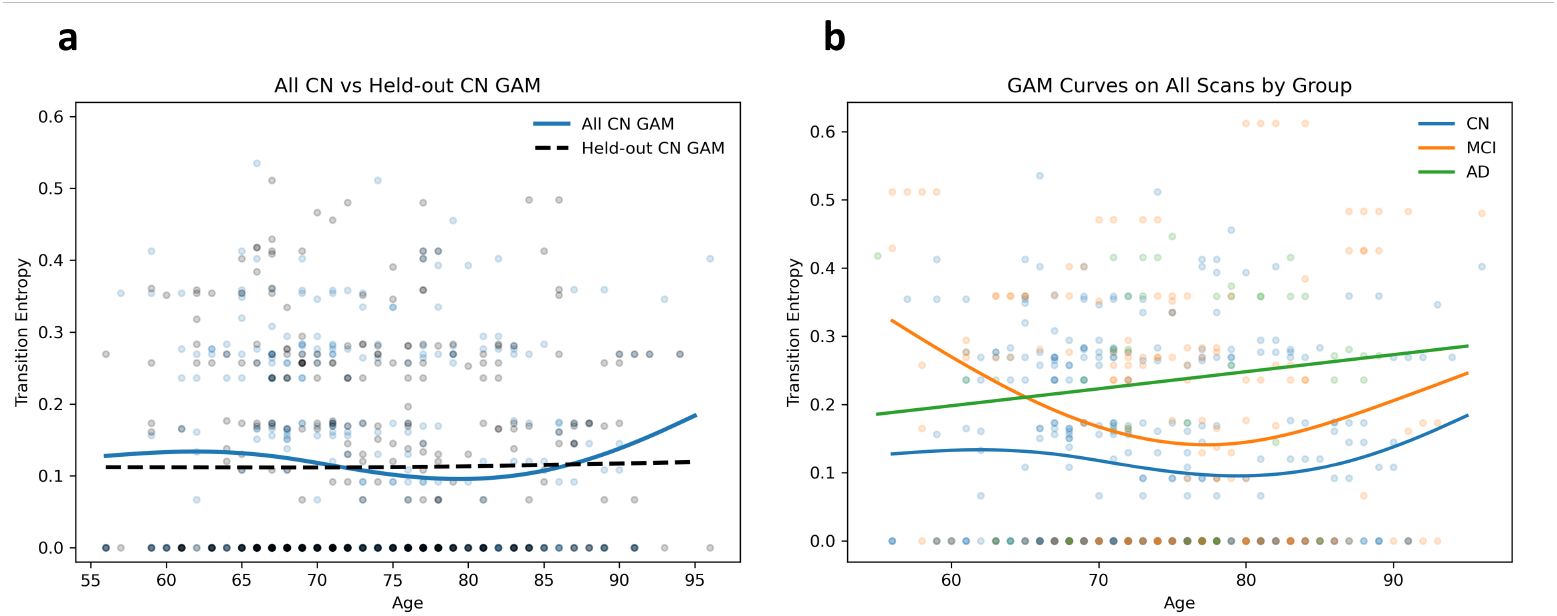
Age-adjusted normative modeling of transition entropy. (a) Generalized additive model (GAM) fitted to cognitively normal (CN) scans and evaluated against held-out CN observations. This panel illustrates the normative age-dependent trajectory used to estimate subject-specific deviation. (b) Group-specific GAM curves for CN, mild cognitive impairment (MCI) and Alzheimer’s disease (AD) scans, shown for descriptive visualization of age-related entropy patterns across diagnostic groups.

Accordingly, we interpret this GAM not as a strong developmental trajectory in itself, but as a normative age-adjusted reference against which disease-related deviations can be quantified.

### 2.2 Alzheimer’s disease shows graded deviation from healthy dynamic aging

Using the normative aging baseline derived from cognitively normal subjects, we next quantified disease-related deviations in brain metastability. For each held-out individual, a deviation score was computed as the difference between the observed transition entropy and the entropy predicted by the CN-trained aging curve at that subject’s age. Figure 2 summarizes the resulting held-out distributions of transition entropy, switching rate, and deviation from the normative CN trajectory across groups. The clearest effect was observed in Alzheimer’s disease (AD), which showed consistently elevated entropy, switching rate, and deviation relative to CN. In particular, mean transition entropy increased from 0.112 in CN to 0.156 in MCI and 0.208 in AD.

**Fig. 2.**
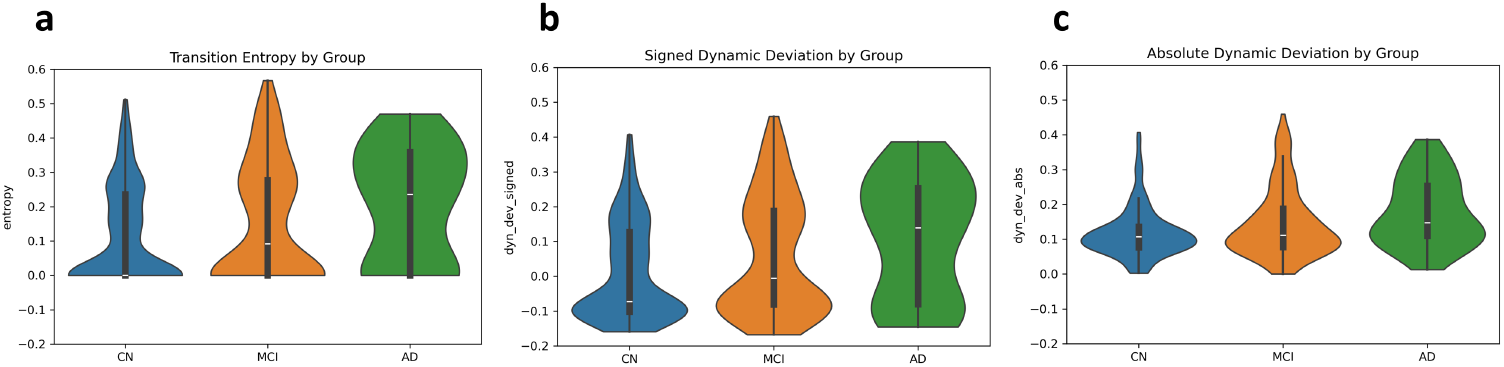
Dynamic disruption across the Alzheimer’s disease continuum. Violin plots show the distribution of (a) transition entropy, (b) signed dynamic deviation and (c) absolute dynamic deviation across cognitively normal (CN), mild cognitive impairment (MCI) and Alzheimer’s disease (AD) groups. Values were obtained from held-out test folds and pooled across cross-validation folds. Transition entropy quantifies the randomness of switching between latent connectivity states, signed deviation captures the direction of departure from the age-adjusted CN normative trajectory, and absolute deviation captures the magnitude of departure irrespective of direction.

Similarly, the mean absolute deviation from the normative CN trajectory was highest in AD (0.178), compared with 0.142 in MCI and 0.107 in CN.

These results indicate that disease-related changes in brain dynamics are not explained by age alone. Rather, AD scans show systematic departures from the age-adjusted CN reference, consistent with abnormal disruption of normative dynamic organization. MCI showed a more intermediate pattern, suggesting that these biomarkers capture an ordered trajectory from healthy aging to MCI and then to AD. However, the substantial overlap between MCI and CN indicates that the proposed biomarkers are most sensitive to AD-related alterations while still reflecting a gradual shift away from healthy dynamics.

### 2.3 Normative deviation provides a compact marker of disease-stage information

To evaluate whether the proposed dynamic measures carried clinically relevant information, we performed subject-level predictive analyses on held-out test folds using three feature settings: transition entropy alone, the absolute deviation from the CN normative trajectory, and the full dynamic feature set, including entropy, switching rate, signed and absolute deviation, fractional occupancy, and dwell time. Prediction was evaluated in both the three-class setting (CN/MCI/AD) and pairwise binary comparisons (Table 1).

**Table 1.** Predictive validation of trajectory-derived dynamic biomarkers across multiclass and pairwise classification tasks.

| Classification | All features |  | Absolute deviation |  | Entropy |  |
| --- | --- | --- | --- | --- | --- | --- |
|  | Accuracy | F1 | Accuracy | F1 | Accuracy | F1 |
| CN/MCI/AD | $0.45 \pm 0.04$ | $0.35 \pm 0.05$ | $0.44 \pm 0.04$ | $0.38 \pm 0.08$ | $0.41 \pm 0.03$ | $0.30 \pm 0.07$ |
| CN/AD | $0.61 \pm 0.06$ | $0.22 \pm 0.04$ | $0.68 \pm 0.07$ | $0.31 \pm 0.14$ | $0.63 \pm 0.09$ | $0.25 \pm 0.11$ |
| CN/MCI | $0.59 \pm 0.05$ | $0.48 \pm 0.05$ | $0.59 \pm 0.04$ | $0.74 \pm 0.09$ | $0.59 \pm 0.04$ | $0.65 \pm 0.07$ |
| AD/MCI | $0.60 \pm 0.07$ | $0.35 \pm 0.14$ | $0.51 \pm 0.16$ | $0.28 \pm 0.16$ | $0.58 \pm 0.10$ | $0.34 \pm 0.13$ |

Values are reported as mean ± standard deviation across five held-out test folds. All features include transition entropy, switching rate, signed deviation, absolute deviation, fractional occupancy and dwell time. CN, cognitively normal; MCI, mild cognitive impairment; AD, Alzheimer’s disease; F1, F1 score.

In the three-class setting, performance remained modest overall, consistent with the substantial overlap between CN and MCI. Nevertheless, the results were informative in two important ways. First, the full feature set achieved the highest accuracy (0.45), indicating that combining multiple complementary summaries of brain dynamics provides the strongest overall discrimination. Second, the single-feature deviation biomarker achieved nearly identical accuracy (0.44) and a higher macro-F1 score (0.38 vs. 0.35), showing that a compact deviation-based summary can retain much of the predictive information contained in the full representation. In addition, entropy alone yielded only moderately lower three-class performance (accuracy = 0.41, macro-F1 = 0.30), indicating that transition entropy by itself remains an informative biomarker. The pairwise analyses further clarified the role of the individual biomarkers. For CN versus AD, Absolute Deviation gave the strongest performance (accuracy = 0.68, F1 = 0.31), exceeding both entropy alone (accuracy = 0.63, F1 = 0.25) and the full feature set (accuracy = 0.61, F1 = 0.22). For CN versus MCI, all three representations achieved the same accuracy (0.59), but Absolute Deviation again yielded the highest F1 score (0.74), compared with 0.65 for entropy and 0.48 for the full feature set. These results suggest that deviation from the CN normative trajectory provides an informative summary of disruption relative to a healthy baseline, even when represented as a single scalar feature. In contrast, for MCI versus AD, the full feature set performed best (accuracy = 0.60, F1 = 0.35), while entropy alone (accuracy = 0.58, F1 = 0.34) outperformed Absolute Deviation (accuracy = 0.51, F1 = 0.28). This suggests that once both groups have already departed from the healthy CN baseline, the raw transition dynamics themselves, particularly entropy, may better capture differences between intermediate and late disease stages than deviation from CN alone.

Taken together, these findings support two complementary conclusions. First, transition entropy and absolute deviation from the CN normative trajectory provide informative single-feature summaries of abnormal brain dynamics. Entropy captures the randomness of switching between latent connectivity states, whereas absolute deviation quantifies the magnitude of departure from age-adjusted healthy dynamics. Second, among these two measures, absolute deviation emerged as a compact candidate biomarker that retained information comparable to the full dynamic feature set, particularly in CN-anchored tasks (CN vs. MCI and CN vs. AD). More broadly, the pattern of results is consistent with an ordered disruption process from CN to MCI to AD: deviation from normative dynamics is modestly detectable in MCI and becomes more pronounced in AD. Thus, these biomarkers are best interpreted not as standalone diagnostic classifiers, but as interpretable biomarkers of disease-related disruption in normative brain-state organization.

## 3 Discussion

Distinguishing pathological neurodegeneration from the broad variability of healthy brain aging remains a central challenge in Alzheimer’s disease research. Here, we addressed this problem by modeling brain dynamics relative to a cognitively normal reference space rather than relying only on group-average differences. Using a CN-trained HMM and an age-adjusted normative trajectory of transition entropy, we show that dynamic brain-state organization exhibits a graded departure from healthy aging across the AD continuum. The clearest alterations were observed in AD, while MCI showed an intermediate pattern with substantial overlap with CN. This pattern suggests that disruption of normative brain dynamics is not merely a static group difference, but may reflect a progressive shift in how the brain occupies and transitions between latent connectivity states. By treating cognitively normal older adults as an age-dependent reference population, this framework places Alzheimer’s disease within the broader challenge of distinguishing normative aging heterogeneity from disease-related deviation.

A central contribution of this work is the use of a normative dynamic-state framework. Instead of training a model to directly discriminate diagnostic groups, we first defined a healthy dynamic reference space from CN subjects and then asked how each subject’s dynamics were expressed relative to that reference. This distinction is important. Standard classification frameworks can identify separable patterns, but they do not directly answer whether an individual’s brain dynamics are abnormal for their age. By contrast, the proposed framework estimates an expected level of transition entropy under healthy aging and quantifies signed and absolute deviations from that expectation. In this sense, the deviation score is not simply another predictive feature; it is an age-adjusted measure of departure from normative brain-state organization.

The biological interpretation of the HMM-derived features is also important. Fractional occupancy reflects how much time the brain spends in particular latent connectivity regimes, dwell time captures the temporal persistence of those regimes, switching rate measures how frequently the brain changes state, and transition entropy quantifies the predictability versus randomness of these transitions. Together, these features summarize complementary aspects of dynamic network organization: state preference, temporal stability, flexibility, and transition structure. In neurodegenerative aging, disruption may therefore appear not only as altered connectivity strength, but also as abnormal temporal coordination among connectivity states. The increased entropy and switching behavior observed in AD are consistent with a less stable or less predictable dynamic regime, suggesting that disease may perturb the temporal organization of large-scale brain networks.

Among the derived biomarkers, absolute deviation from the CN normative trajectory emerged as a particularly compact and interpretable measure. Despite being a single scalar feature, it remained competitive with the full dynamic feature set in predictive validation and performed especially well in CN-anchored comparisons. This supports the idea that the magnitude of departure from healthy age-adjusted dynamics captures a meaningful axis of disease-related disruption. At the same time, entropy itself remained informative, particularly for distinctions involving later disease stages. This suggests that deviation from the healthy reference and raw transition randomness capture related but non-identical aspects of neurodegenerative change: deviation quantifies distance from normative aging, whereas entropy reflects the internal organization of state switching itself.

The predictive analyses provide further support for the clinical relevance of these biomarkers, but also highlight their intended role. Three-class prediction across CN, MCI, and AD remained modest, reflecting the biological and clinical overlap among diagnostic categories, especially between CN and MCI. Pairwise analyses showed clearer patterns, with absolute deviation performing strongly in CN-anchored tasks and entropy or the broader dynamic feature set contributing to later-stage distinctions. These findings suggest that the proposed features are better interpreted as progression-sensitive biomarkers rather than standalone diagnostic classifiers. Their value lies less in replacing clinical diagnosis and more in providing an interpretable measure of how far an individual’s brain dynamics have departed from healthy aging. This interpretation is consistent with the conceptual organization of the AD continuum. MCI is clinically heterogeneous: some individuals remain stable, some revert, and some progress to AD. Therefore, substantial overlap between CN and MCI is not necessarily a failure of the model, but may reflect the biological heterogeneity of the MCI category itself. In contrast, AD showed the most pronounced shift away from the normative CN trajectory, suggesting that disruption of brain-state dynamics becomes increasingly evident with more advanced pathology. A particularly important next step will be to evaluate whether these dynamic deviations predict longitudinal conversion from CN to MCI or from MCI to AD. Such an analysis would directly test whether the proposed biomarkers capture progression rather than cross-sectional diagnostic status alone.

Several methodological choices strengthen the interpretability of the proposed framework. First, PCA, HMM training, and normative modeling were performed within cross-validation folds using training CN data only, reducing the risk that disease or held-out test information influenced the learned reference space. Second, final biomarker distributions and prediction metrics were based on held-out outputs pooled across folds, so each subject contributed to the reported results only when evaluated out of sample. Third, the HMM was trained on CN data to define a common dynamic state space, allowing MCI and AD subjects to be evaluated according to how they navigate a healthy reference system rather than by learning disease-specific states. This design directly supports the interpretation of disease effects as deviations from normative dynamics.

There are also important limitations. The analysis is based on resting-state fMRI, which is sensitive to preprocessing choices, motion, sampling variability, and the choice of parcellation. The ADNI cohort is multi-site, and although standard preprocessing and nuisance regression were applied, residual effects of motion, scanner/site differences, and acquisition variability may influence resting-state dynamic connectivity estimates. The AD group was relatively small compared with the CN and MCI groups, which may affect the stability of group estimates and pairwise prediction metrics. Sliding-window connectivity introduces additional assumptions about window length and temporal resolution, and although the HMM provides a structured model of state transitions, the number of states remains a modeling choice that should be assessed through sensitivity analyses.

Although the present study evaluated multiple HMM-derived dynamic summaries, the normative component of the framework was centered on transition entropy because it provided the clearest and most interpretable age-adjusted trajectory. A key next step is to develop a longitudinal trajectory-based framework for modeling how brain-state dynamics change within individuals as they transition across clinical stages of Alzheimer’s disease. By tracking deviation from normative aging patterns over time, such an approach could test whether dynamic biomarkers capture within-subject progression from CN to MCI or from MCI to AD, rather than cross-sectional disease-stage differences alone.

In summary, this study reframes dynamic functional connectivity in Alzheimer’s disease as a problem of deviation from normative brain-state organization. By anchoring HMM-derived dynamics to an age-adjusted CN reference, we show that neurodegenerative pathology is associated with progressive disruption of latent brain-state transitions. The resulting biomarkers, especially transition entropy and absolute deviation from the normative trajectory, provide interpretable and compact measures of abnormal dynamic organization. Rather than serving solely as diagnostic classifiers, these measures may offer a principled way to quantify progression-sensitive disruption of brain dynamics across the Alzheimer’s disease continuum.

## 4 Online Methods

### 4.1 Data and Cohort Definition

Resting-state functional MRI (rs-fMRI) data are obtained from the Alzheimer’s Disease Neuroimaging Initiative (ADNI), including cognitively normal (CN), mild cognitive impairment (MCI), and Alzheimer’s disease (AD) participants. Each subject contributed one or more rs-fMRI scans and a T1-weighted structural image. Diagnostic labels are harmonized across metadata files and encoded as an ordinal disease stage variable (CN = 0, MCI = 1, AD = 2) for downstream analyses. Demographic information, including chronological age, was used only for statistical association analyses.

The final cohort comprised 357 participants (238 CN, 94 MCI, 25 AD) contributing a total of 585 resting-state fMRI scans, with some subjects contributing multiple longitudinal sessions (Table 2). Groups did not differ significantly in age at the time of first scan (one-way ANOVA, F(2,354)=2.77, p=0.064), supporting the use of a normative age-adjusted modeling framework. Scans were acquired between 2017 and 2023 across multiple ADNI sites.

**Table 2.** Demographic and scan characteristics of the ADNI cohort.

| Characteristic | CN | MCI | AD |
| --- | --- | --- | --- |
| Subjects, $n$ | 238 | 94 | 25 |
| Scans, $n$ | 373 | 172 | 40 |
| Scans per subject, mean (range) | 1.6 (1–4) | 1.8 (1–5) | 1.6 (1–3) |
| Age at first scan, mean ± s.d. | 72.3 ± 7.8 | 74.1 ± 8.5 | 75.1 ± 8.2 |
| Age range, years | 56–96 | 56–96 | 55–88 |
| Scan years | 2017–2023 |  |  |

Age statistics were computed from each subject’s first available scan. Groups did not differ significantly in age by one-way ANOVA, $F(2,354) = 2.77$ , $p = 0.064$ . CN, cognitively normal; MCI, mild cognitive impairment; AD, Alzheimer’s disease; s.d., standard deviation.

### 4.2 Data Conversion and Preprocessing

Raw MRI data were originally acquired in DICOM format. For each subject, structural (T1-weighted) and resting-state functional images were converted to NIfTI format before preprocessing. Converted files were organized by subject and modality and served as input to the preprocessing pipeline.

Preprocessing was performed using SPM12 in combination with the CONN toolbox through batch processing to ensure reproducibility. For each subject, one T1-weighted structural image and one resting-state fMRI run were included. Functional images were acquired with a repetition time (TR) of 3 s.

The preprocessing pipeline consisted of the following steps, applied sequentially:

1. Functional centering, 2. Slice-timing correction using ascending slice acquisition order, 3. Motion correction via realignment and unwarping, 4. Coregistration of functional images to the structural T1 image, 5. Structural segmentation into gray matter, white matter, and cerebrospinal fluid, 6. Spatial normalization to MNI space using segmentation-derived deformation fields, and 7. Spatial smoothing with a Gaussian kernel (FWHM = 6 mm).

Temporal denoising is performed within CONN, including band-pass filtering in the range 0.008–0.09 Hz and standard nuisance regression to reduce non-neural confounds. Preprocessing outputs included normalized functional images, ROI-level time series, and quality-control measures.

Brain regions of interest were defined using the Craddock atlas, comprising 840 parcels. The atlas was treated as a multi-label parcellation in standard space. Following preprocessing, mean BOLD time series were extracted for each ROI and subject, yielding an ROI-by-time matrix that served as the basis for all subsequent analyses.

### 4.3 Dynamic Functional Connectivity

To capture temporal variability in functional connectivity, dynamic functional connectivity (dFC) matrices were computed using a sliding-window approach[8, 10, 11]. A window length of 40 TRs (120 seconds at TR = 3 s) was selected to balance temporal resolution against the statistical reliability of windowed correlation estimates[22, 23]. Let **X**^(*s*)^ ∈ ℝ^*T* ×*R*^ denote the preprocessed resting-state fMRI time series for subject *s*, where *T* is the number of time points and *R* is the number of regions of interest (ROIs).

Using a sliding window of length *L* and step size Δ, windowed segments are defined as

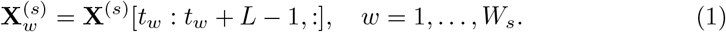

For each window, the dynamic functional connectivity (dFC) matrix is computed using Pearson correlation and Fisher *z*-transformation on the columns of 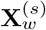 as follows:

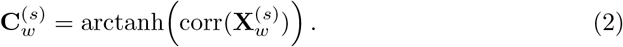

Although sliding-window dFC has known methodological limitations, including sensitivity to window length and potential sampling variability, it remains a widely used framework for characterizing time-varying functional connectivity [22, 24, 25].

### 4.4 Feature Vectorization and Dimensionality Reduction

For each windowed functional connectivity matrix, the upper-triangular elements were extracted and vectorized to form a high-dimensional feature vector. This representation preserved all unique pairwise connectivity values while avoiding redundancy from matrix symmetry. The Fisher Z-transform is also applied to correlation values (after clipping) to stabilize variance and make them more Gaussian-like before any dimensionality reduction step.

Given the high dimensionality of window-level connectivity vectors, dimensionality reduction was performed using principal component analysis (PCA). To avoid data leakage and preserve a strictly normative framework, PCA was estimated within each cross-validation fold using only the window-level FC vectors from the training cognitively normal (CN) subjects. The learned PCA transformation was then applied unchanged to all subjects in that fold, including held-out CN, mild cognitive impairment (MCI), and Alzheimer’s disease (AD) subjects. This procedure ensured that all subjects were represented in a common low-dimensional feature space defined only by healthy training data.

The resulting reduced-dimensional window representations were then used as the observations for the HMM, allowing the model to characterize dynamic connectivity states in a CN-defined reference space. Figure 3 summarizes the full normative dynamic-connectivity framework used in this study.

**Fig. 3.**
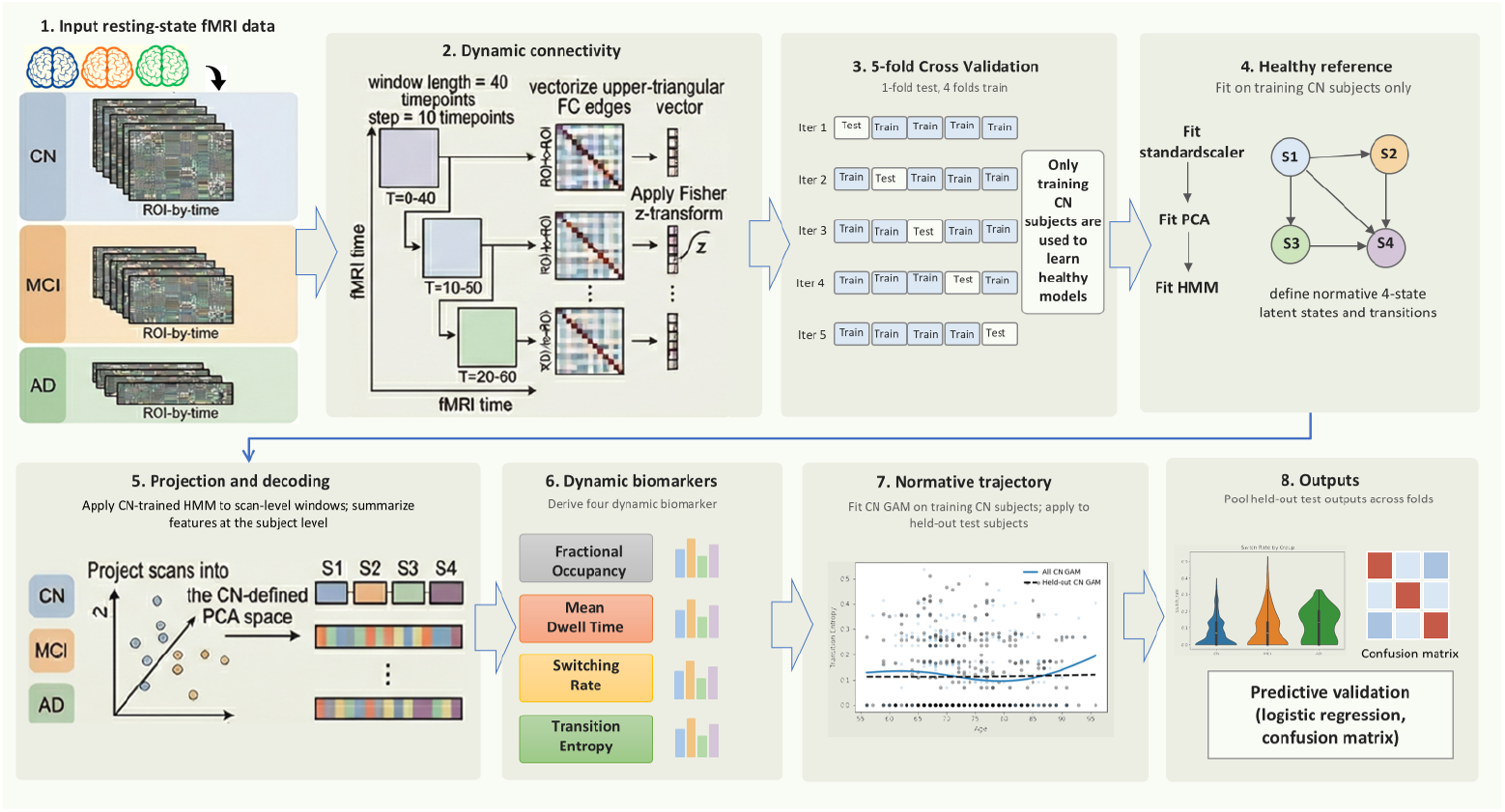
Normative dynamic-connectivity pipeline for modeling brain-state disruption across aging and Alzheimer’s disease. Resting-state fMRI time series from cognitively normal (CN), mild cognitive impairment (MCI), and Alzheimer’s disease (AD) subjects were converted into sliding-window functional connectivity matrices (window length = 40 time points, step = 10). Upper-triangular FC edges were vectorized and Fisher z-transformed. In stratified 5-fold cross-validation, one fold was held out for testing, and the remaining four folds were used for training, while only training CN subjects were used to define the healthy reference space. Specifically, a scaler, PCA model, and 4-state HMM were fit on training CN data only, yielding a CN-defined latent dynamic state space. The trained PCA and HMM were then applied to all subjects in the fold to decode state sequences, from which dynamic biomarkers were derived, including fractional occupancy, mean dwell time, switching rate, and transition entropy. A CN-only generalized additive model (GAM) was then fitted to model the normative age-dependent entropy trajectory, and signed and absolute deviations from this trajectory were computed for held-out test subjects. Final held-out outputs were pooled across folds for group-level biomarker analysis and predictive validation.

### 4.5 Hidden Markov Model of Dynamic Connectivity

Dynamic connectivity sequences were modeled using a Hidden Markov Model (HMM) [12, 16] with *K*_*s*_ latent states. Let *q*_*t*_ ∈ {1, … , *K*_*s*_} denote the latent state at window *t*. State transitions follow a first-order Markov process:

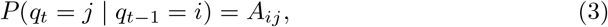

where 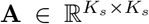 is the transition matrix. The observations consisted of PCA-reduced connectivity vectors, and state-specific emission distributions were modeled as Gaussian distributions with diagonal covariance.

To define a normative healthy state space, the HMM was trained using CN subjects only. The resulting CN-trained model was then applied to all subjects to infer posterior state probabilities and the most likely latent state sequence for each time window.

To characterize individual brain dynamics, window-level state assignments were summarized into subject-level metrics. These included: 1)*Fractional Occupancy*, defined as the average posterior probability of occupying each latent state across windows, 2)*Mean Dwell Time*, defined as the average duration of consecutive visits to a given state, 3)*Transition Entropy*, quantifying the predictability versus randomness of state switching, and 4)*Switching Rate*, defined as the proportion of state changes across consecutive windows.

These features provide complementary summaries of brain dynamics. Fractional occupancy reflects how much time the brain spends in different connectivity regimes, mean dwell time captures the temporal stability or persistence of those regimes, transition entropy quantifies the predictability versus variability of switching between states, and switching rate summarizes the overall volatility of state transitions. In the context of neurodegenerative aging, alterations in these measures may reflect disrupted coordination, reduced flexibility, or abnormal stabilization of large-scale brain network dynamics.

### 4.6 Normative Modeling of Dynamic Deviations

To quantify disease-related deviation from healthy aging dynamics, age-adjusted normative models were estimated using only cognitively normal (CN) subjects. Within each cross-validation fold, a generalized additive model (GAM) [18, 26, 27] was fitted using the training CN fMRI scans, with age as a smooth predictor of transition entropy. Following standard GAM notation[26], the normative model takes the form:

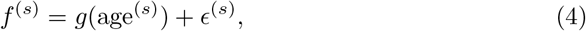

where *f* (*s*) denotes the transition entropy of scan *s, g*(·) is a smooth nonlinear function of age, and *ϵ*(*s*) is the residual term.

The fitted CN-based GAM was then applied to all held-out scans in that fold, including CN, MCI, and AD, to estimate the entropy expected under normative healthy aging at the corresponding age. For each scan, first, a signed deviation score was computed as the difference between the observed entropy and the age-adjusted CN prediction:

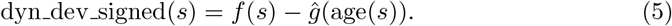

Absolute deviation was then defined as:

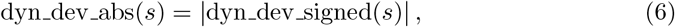

which quantifies the magnitude of departure from the normative CN trajectory irrespective of direction. These deviation measures were used as biomarkers of abnormal dynamic brain organization relative to age-adjusted healthy dynamics.

### 4.7 Prediction of Dementia Stage from Dynamic Features

To evaluate whether the extracted dynamic biomarkers carried disease-stage information, we performed subject-level prediction analyses using multinomial logistic regression. The primary evaluation used stratified 5-fold cross-validation[28], such that in each iteration one fold served as the held-out test set and the remaining four folds were used for training. Cross-validation splits were performed at the subject level, such that all scans from a given participant were assigned to the same fold. This prevented scans from the same individual from appearing in both training and held-out test sets. To avoid data leakage, all preprocessing and modeling steps, including standardization, PCA, HMM training, and CN-based normative modeling, were performed using the training data only within each fold.

We evaluated three feature settings: (i) transition entropy alone, (ii) the absolute deviation from the CN normative trajectory (dyn dev abs) alone, and (iii) the full dynamic feature set consisting of transition entropy, switching rate, signed and absolute deviation, fractional occupancy, and dwell time. All predictor variables were standardized within each training fold before classifier fitting.

Prediction performance was assessed in both the three-class setting (CN /MCI /AD) and the pairwise binary settings (CN versus MCI, CN versus AD, and MCI versus AD). Performance was summarized using accuracy, F1 score, and confusion matrices.

It is important to note that some pairwise tasks, particularly CN versus AD, were class-imbalanced, with substantially fewer AD scans than CN scans. As a result, accuracy and F1 did not always align: relatively high accuracy could still coincide with lower F1 when the minority class was not detected as reliably. For this reason, we interpret the pairwise results using both metrics rather than accuracy alone[29, 30].

## Data availability

Data used in the preparation of this article were obtained from the Alzheimer’s Disease Neuroimaging Initiative (ADNI) database (adni.loni.usc.edu). As such, the investigators within the ADNI contributed to the design and implementation of ADNI and/or provided data but did not participate in the analysis or writing of this report. A complete listing of ADNI investigators can be found at: http://adni.loni.usc.edu/wp-content/uploads/how-to-apply/ADNI-Acknowledgement-List.pdf

## Code availability

The source code used in this research is openly available on GitHub at https://github.com/monirt/Normative-trajectory-modeling-of-brain-dynamics-across-aging-and-AD.

## Acknowledgements

The authors used ChatGPT solely for language editing and clarity improvements. All scientific content, analyses, interpretations, and conclusions are entirely the authors’ own.

Data collection and sharing for the Alzheimer’s Disease Neuroimaging Initiative (ADNI) is funded by the National Institute on Aging (National Institutes of Health Grant U19AG024904). The grantee organization is the Northern California Institute for Research and Education. In the past, ADNI has also received funding from the National Institute of Biomedical Imaging and Bioengineering, the Canadian Institutes of Health Research, and private sector contributions through the Foundation for the National Institutes of Health (FNIH) including generous contributions from the following: AbbVie, Alzheimer’s Association; Alzheimer’s Drug Discovery Foundation; Araclon Biotech; BioClinica, Inc.; Biogen; Bristol-Myers Squibb Company; CereSpir, Inc.; Cogstate; Eisai Inc.; Elan Pharmaceuticals, Inc.; Eli Lilly and Company; EuroImmun; F. Hoffmann-La Roche Ltd and its affiliated company Genentech, Inc.; Fujirebio; GE Healthcare; IXICO Ltd.; Janssen Alzheimer Immunotherapy Research & Development, LLC.; Johnson & Johnson Pharmaceutical Research & Development LLC.; Lumosity; Lundbeck; Merck & Co., Inc.; Meso Scale Diagnostics, LLC.; NeuroRx Research; Neurotrack Technologies; Novartis Pharmaceuticals Corporation; Pfizer Inc.; Piramal Imaging; Servier; Takeda Pharmaceutical Company; and Transition Therapeutics.

## Author contributions

MT conceptualized the study, performed statistical analysis, drafted the manuscript, and led the revisions in response to reviewer feedback. VR provided critical revisions, methodological guidance, and editorial support throughout the process. All authors contributed to refining the manuscript and approved the final version.

## Additional information

Supplementary information is available for this paper.

