## Supplementary Information for "Normative brain-state trajectories reveal deviation from healthy aging in AD"

Monir Taimouri and Vikram Ravindra

**Correspondence:** Monir Taimouri, Department of Computer Science, University of Cincinnati.

**This PDF file includes:**

- Supplementary Notes 1–3
- Supplementary Tables 1–4

### Supplementary Note 1. HMM state-number sensitivity analysis

To assess whether the main findings depend on the number of HMM latent states, the pipeline was repeated with $K = 5$ and $K = 6$. The HMM was trained on unique CN subjects ($N = 238$) and evaluated on unique held-out subjects (CN = 238, MCI = 94, AD = 25). Group differences were assessed with the Kruskal–Wallis test.

The group effect on transition entropy was statistically significant across all three configurations: $K = 4$ ($H = 20.26$, $p < 0.001$), $K = 5$ ($H = 12.00$, $p = 0.002$), and $K = 6$ ($H = 8.64$, $p = 0.013$). The ordered pattern CN < MCI < AD was preserved for $K = 4$ and $K = 6$. For $K = 5$, the pattern was preserved in entropy (CN = 0.084, MCI = 0.120, AD = 0.128), although the MCI–AD gap was smaller.

**Table 1. Kruskal–Wallis test results and group mean transition entropy for $K = 4$, $K = 5$ and $K = 6$ HMM states.** Values are not directly comparable in scale across rows, but the direction of the group effect is preserved.

| $K$ | CN | MCI | AD | KW $H$ | $p$ value |
|---|---|---|---|---|---|
| 4 | 0.112 | 0.156 | 0.208 | 20.26 | < 0.001 |
| 5 | 0.084 | 0.120 | 0.128 | 12.00 | 0.002 |
| 6 | 0.107 | 0.115 | 0.175 | 8.64 | 0.013 |

**Table 2. Group mean dynamic biomarkers for $K = 5$ and $K = 6$.** The ordered pattern is preserved across features, most clearly for $K = 6$.

| $K$ | Group | Entropy | Switching rate | Signed deviation | Absolute deviation |
|---|---|---|---|---|---|
| 5 | CN | 0.084 | 0.058 | 0.006 | 0.100 |
| | MCI | 0.120 | 0.088 | 0.044 | 0.123 |
| | AD | 0.128 | 0.088 | 0.051 | 0.126 |
| 6 | CN | 0.107 | 0.081 | 0.025 | 0.098 |
| | MCI | 0.115 | 0.096 | 0.036 | 0.102 |
| | AD | 0.175 | 0.137 | 0.099 | 0.136 |

### Supplementary Note 2. Within-subject PCA centering and group discrimination

Within-subject centering in PCA space removes each subject's static mean connectivity profile before HMM fitting, retaining only within-subject temporal fluctuations. Although this can improve sensitivity to dynamic variation, it may also remove between-subject variance that carries diagnostically relevant information. We compared group discrimination with and without centering using $K = 4$.

Without centering, the Kruskal–Wallis test yielded $H = 20.26$ ($p < 0.001$), with mean transition entropy of 0.112 for CN, 0.156 for MCI and 0.208 for AD. With centering applied, the statistic decreased to $H = 6.76$ ($p = 0.034$), and entropy values were compressed to approximately 0.083 for CN, 0.088 for MCI and 0.131 for AD. On this basis, the main analysis was conducted without within-subject centering.

**Table 3. Transition entropy group statistics with and without within-subject PCA centering.** Results are shown for $K = 4$ using unique subjects ($N = 357$).

| Centering | CN | MCI | AD | KW $H$ | $p$ value |
|---|---|---|---|---|---|
| No, main analysis | 0.112 | 0.156 | 0.208 | 20.26 | $< 0.001$ |
| Yes | 0.083 | 0.088 | 0.131 | 6.76 | 0.034 |

### Supplementary Note 3. Region-of-interest validation

The main analysis used all 840 Craddock atlas parcels. To evaluate whether the biomarker reflects local medial temporal lobe degeneration or broader network-level disruption, we repeated the HMM analysis on two anatomically restricted parcel subsets.

#### Hippocampal and entorhinal region of interest

Hippocampal parcels were identified by overlapping the Harvard–Oxford subcortical atlas, including the left and right hippocampus, with the Craddock 840-parcel atlas, yielding 24 parcels. An entorhinal cortex proxy was defined using the parahippocampal gyrus, anterior division, yielding 24 parcels. Their union comprised 39 unique parcels.

The Kruskal–Wallis test was significant ($H = 13.03$, $p = 0.001$). However, group means were nearly identical between MCI (0.223) and AD (0.222), with AD not showing higher entropy than MCI. This contrasts with the whole-brain result, indicating that the biomarker is not driven by hippocampal dynamics alone.

#### Broader AD-relevant network

A broader AD-relevant network was defined comprising the bilateral hippocampus, bilateral amygdala, parahippocampal gyrus, angular gyrus, frontal medial cortex, posterior cingulate gyrus and precuneus, mapped to 99 Craddock parcels.

The HMM with $K = 4$ yielded $H = 16.13$ ($p < 0.001$), with group means of 0.205 for CN, 0.217 for MCI and 0.240 for AD, preserving the ordered pattern but with smaller separation than the whole-brain analysis.

**Table 4. Whole-brain, AD-network and hippocampal/entorhinal ROI comparison.** The hippocampal/entorhinal ROI analysis showed nearly identical MCI and AD entropy, suggesting that the whole-brain biomarker is unlikely to be reducible to local hippocampal/entorhinal dynamics alone.

| Analysis | Parcels | CN | MCI | AD | KW $H$ | $p$ value |
|---|---|---|---|---|---|---|
| Whole brain, main analysis | 840 | 0.112 | 0.156 | 0.208 | 20.26 | $< 0.001$ |
| AD network | 99 | 0.205 | 0.217 | 0.240 | 16.13 | $< 0.001$ |
| Hippocampal/entorhinal ROI | 39 | 0.210 | 0.223 | 0.222 | 13.03 | 0.001 |
